# Enhancing lysosome function via mTOR/TFEB activation reduces lipofuscin-like granules in early Age-related Macular Degeneration

**DOI:** 10.1101/2024.09.17.613413

**Authors:** Ana S Falcão, Mafalda Lopes-da-Silva, Pedro Antas, Cristina Escrevente, Margarida Pedro, Rita Coelho, Inês S Ferreira, Inês P Santos, Thomas Ciossek, Paul Nicklin, Sandra Tenreiro, Miguel C Seabra

## Abstract

Age-related macular degeneration (AMD) is the most common blinding disease in the western world and is currently incurable. Although the exact causes of AMD are not clear, the primary origin of pathology appears to be in the retinal pigment epithelium (RPE). RPE is responsible for the daily digestion of photoreceptor outer segments (POS), which imposes a heavy continuous burden on the lysosomal network. POS feeding assay *in vitro* suggested that the accumulation of autofluorescence granules (AFG), similar to lipofuscin *in vivo,* derives from lysosomal dysfunction. Here we show that synchronous phagocytosis of POS leads to early transient mTOR activation followed by inhibition in late phagosome maturation. One of its substrates, the transcription factor EB (TFEB) increases during phagosome maturation albeit mostly in its inactive phosphorylated form. We questioned whether induction of the mTOR/TFEB axis could improve digestion of POS and hence reduce AFG load. Treatment of POS-fed cells with rapamycin, an mTORC1 inhibitor after the appearance of AFG results in 30% reduction of AFG load. This effect is dependent on active lysosomal enzymes and induction of active dephosphorylated TFEB with consequent activation of GADD34 and lysosomal biogenesis. As a proof of concept, we show that overexpressing a constitutively active form of unphosphorylated TFEB dramatically reduces POS-dependent AFG accumulation. Overall, this study suggests that viral or pharmacological approaches activating the mTOR/TFEB axis in the RPE could be beneficial as cell-protective treatment of early/intermediate cases of AMD, acting to delay progression of the disease.

## INTRODUCTION

Age-related macular degeneration (AMD) is the leading cause of visual impairment in the Western world, characterized by loss of vision in the macular central area of the retina. With the increase of the elderly population during the last decades, especially in developed countries, AMD is becoming a major public health concern. Currently, there are no effective treatments for the common forms of AMD namely early, intermediate, or late-stage “dry” AMD, also known as geographic atrophy (*1, 2*).

Although the exact causes of AMD are not clear, the primary origin of pathology appears to be in the retinal pigment epithelium (RPE). Findings suggest that lysosomal dysfunction in RPE cells, cellular senescence, and abnormal immune-inflammatory responses are all involved in AMD pathogenesis and promote its progression. These factors interact with each other, causing lipofuscin deposition, drusen formation, RPE injury or atrophy, which can lead to photoreceptor cell damage, choroid degeneration, and ultimately, loss of vision (*3*). In fact, the RPE is responsible for the daily digestion of photoreceptor outer segments (POS), a process essential for sustained photoreceptor function which requires the regular recycling of visual cycle components (*4*). However, POS digestion by RPE cells imposes a heavy continuous burden on the lysosomal network of these post-mitotic cells. In aged RPE cells, lipofuscin accumulation due to undegraded phagocytosed POS results in the accumulation of autofluorescent (AF) material (*5*). RPE cells normally cope with accumulated lipofuscin but in intermediate AMD, excess lipofuscin is a sign of RPE stress (*6*). We recently described an *in vitro* model system whereby feeding RPE cells with a single pulse of porcine POS leads to the accumulation of autofluorescent granules (AFG), similar to lipofuscin *in vivo* (*7*). We observed that lysosome function, in particular lysosome-phagosome fusion is required for AFG formation but that impairment of lysosome catalytic activity, particularly Cathepsin D (CTSD), enhanced AFG accumulation. Therefore, dysfunctional lysosomal activity leads to reduced POS degradation and increased AFG accumulation.

Autophagic dysregulation has been suggested to occur in neurodegenerative diseases (*8*), including AMD (*9, 10*). Correspondingly, impaired autophagy has been observed in RPE of AMD donor tissue and multiple models of inherited retinal degenerations (*4*). Autophagy proteins, autophagosomes, and autophagy flux were significantly reduced in tissue from human donor AMD eyes and animal models of AMD (*11*). Moreover, RPE cultures established from AMD donors exhibited several features of disrupted autophagy, such as swollen autolysosomes and impaired autophagic flux (*12*). AMD donor RPE cultures also accumulated lipid droplets, glycogen granules, and breakdown of mitochondria, indicating an impairment of cellular clearance pathways. Altogether, evidence suggests that RPE lysosomal dysfunction could be an important feature in early and intermediate AMD. However, many questions remain unanswered, such as the cross-talk between POS phagocytosis with lysosome biogenesis and/or autophagy machinery, as well as which specific steps of POS degradation and which lysosome-related proteins are impaired in AMD.

The transcription factor EB (TFEB) is a member of the MIT-TFE helix–loop– helix leucine-zipper (bHLH-Zip) family of transcription factors that includes other members such as micropthalmia-associated transcription factor (MITF), TFE3, and TFEC (*13*). TFEB is regulated by the mammalian target of rapamycin (mTOR) pathway, *i.e.*, under anabolic conditions when there are nutrients available, mTOR complex 1(mTORC1) is activated at the lysosomal membrane leading to TFEB phosphorylation and thus its inactivation in the cytosol (*14*). Conversely, under stress or starvation causing catabolic conditions, mTORC1 is inhibited following release from the lysosome membrane. Unphosphorylated TFEB is translocated to the nucleus resulting in transcriptional activation of its target genes. TFEB is described as a major controller of lysosomal biogenesis and autophagy by positively regulating genes belonging to the Coordinated Lysosomal Expression and Regulation (CLEAR) network (*14, 15*) .

As a master modulator of intracellular clearance pathways, TFEB has been proposed as an important therapeutic target in diseases characterized by accumulation of toxic aggregates, including lysosomal storage disorders (*16, 17*), neurodegenerative diseases (*18*) and vision diseases (*19*). In cells that are highly involved in lysosome-mediated degradation processes, such as RPE, mTORC1 appears to be the primary regulator of TFEB activity. Furthermore, TFEB overexpression increases transcription of autophagy and lysosomal genes, thus leading to the accumulation of autophagosomes (*14, 20*). Therefore as lysosomal degradative efficiency declines with age and dysfunction in cellular clearance has been implicated in AMD, we hypothesized that TFEB may be a potential therapeutic target for this disease.

The mTOR signalling pathway and/or the autophagic process may also be a potential therapeutic target for this disease. In fact, mTOR inhibitors and autophagy inducers have been recently considered as potential therapeutic strategies for AMD (*21, 22*). Rapamycin, a lead mTOR inhibitor, is an attractive option with a favourable safety profile and sustained ocular pharmacokinetics, although clinical trials in advanced AMD showed limited efficacy results and some adverse effects (reviewed in (*21*)). Thus, understanding the mTOR/TFEB pathway and the mechanisms involved during the phagocytosis and clearance of POS could reveal new therapeutic targets to avoid formation and/or decrease the accumulation of lipofuscin in AMD. Here we test these hypothesis and our results suggest that the TFEB/mTOR axis could be a potential therapeutic approach to reduce AFG accumulation and delay AMD progression.

## RESULTS

### TFEB gene expression is induced 72h after POS feeding

In our previous work, we presented a new cellular model of AFG formation by feeding RPE cells with POS (*7*). In the current study using the same *in vitro* model, we focused on exploring the molecular mechanisms of POS phagocytosis and clearance as a way to reveal potential therapeutic windows and/or targets to decrease the AF load of these cells. To explore these pathways, in vitro differentiated monolayers of RPE (ARPE-19 and hfRPE) cells were pulsed with POS during 4h, followed by a chase period of up to 7 days. We then analysed the expression of TFEB and other lysosomal biogenesis-related genes and proteins (Fig 1).

**Fig. 1.**
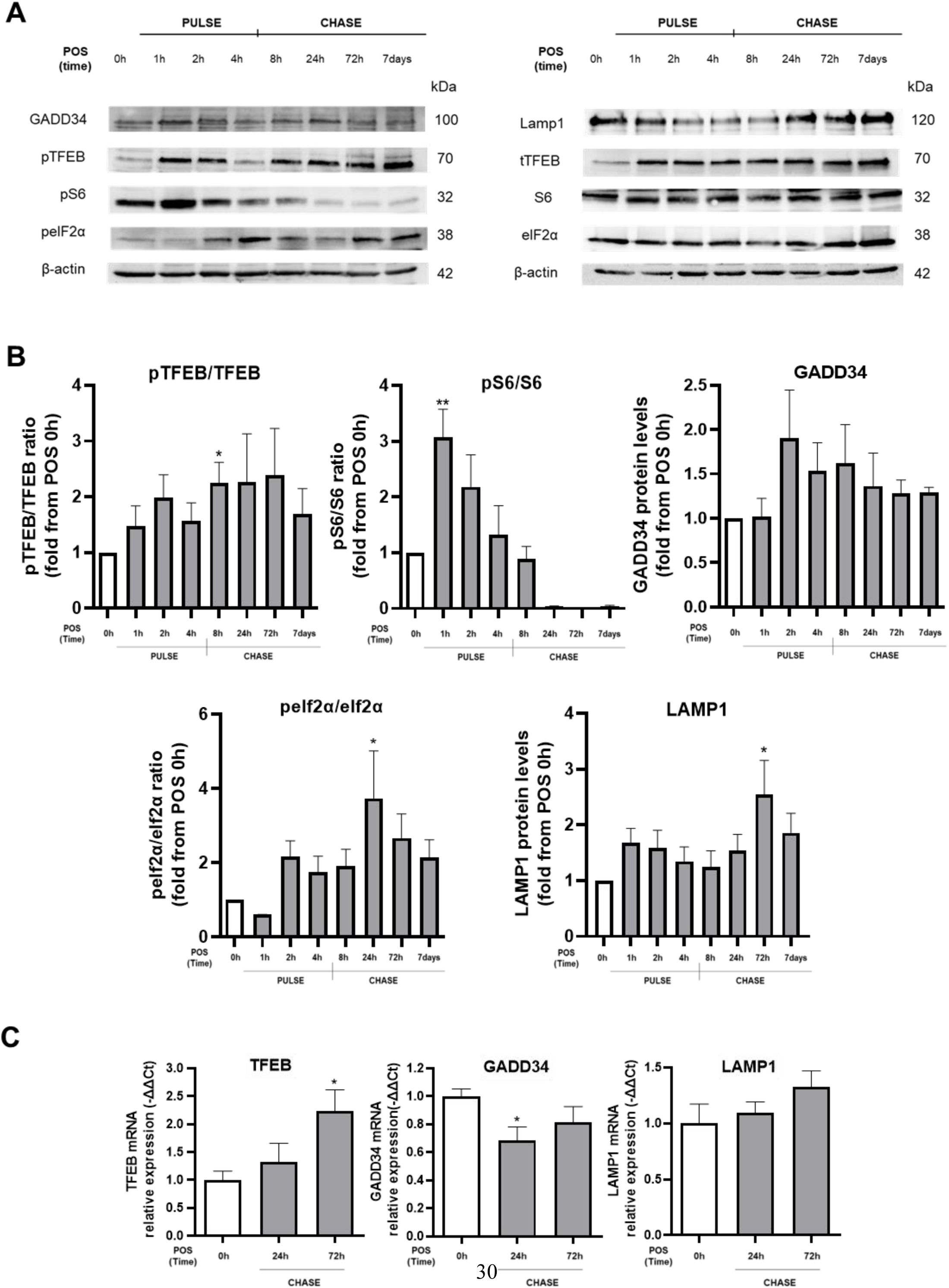
Mechanisms of POS phagocytosis and clearance during the pulse and chase periods. ARPE-19 cells were pulsed during 4h with 200 μg/mL POS and chased for 7 days. (A) Evaluation of pTFEB, total TFEB, pS6, total S6, GADD34, peIf2α, total peIf2α, Lamp1 and β-actin protein levels at the indicated time points during pulse and chase periods, assessed by western blot. Representative blots of one experiment are shown. (B) Densitometric analysis of the immunodetection of the indicated proteins was performed as described in Materials and Methods and the ratio of pTFEB/TFEB, pS6/S6 and peIf2α/eIf2α were calculated, using β-actin as loading control. (C) mRNA expression levels of TFEB, GADD34 and LAMP1 at the indicated time points, analysed by qPCR using HRPT1 as housekeeping gene. All data are shown as mean ± SEM of at least 3 independent experiments (n≥3). A one-way ANOVA followed by multiple comparisons Dunnet post hoc correction was used for statistical analysis, except for the ratio of pTFEB/TFEB and GADD34 mRNA relative expression that was analysed using one-way ANOVA Kruskal Wallis test. *P<0.05 and **P<0.01 vs. no POS (white bars, 0h).

We observed that 1h immediately after the POS pulse there is a significant increase in the pS6 (S6-kinase beta-1) /S6 ratio (Fig 1A and B), which then decreases during the chase period. This ribosomal protein is a downstream target of mTOR, as it is phosphorylated and activated by mTORC1 (*21*). This result suggests that during the pulse with POS there is an early activation of mTOR to aid in the proper POS phagocytosis and degradation, as described in mouse RPE *in vivo* (*23*). Furthermore, we detected a significant increase in TFEB gene expression in the 72h chase period (Fig 1C). However, TFEB levels trend towards an increased pTFEB/TFEB ratio in the chase period (although only significant at the 8h timepoint), suggesting that a significant fraction of TFEB (pTFEB) could be inactive (Fig 1A and B). Interestingly, this increase in pTFEB is accompanied by a significant decrease in GADD34 gene expression (24h chase period) (Fig. 1C). GADD34 is a component of the protein phosphatase 1 complex recently described as an early and direct TFEB target, which functions as an enabler of lysosomal biogenesis and autophagy by dephosphorylating eIF2α (*24*). As expected, lower GADD34 gene expression levels during this period led to a significant increase of its substrate, the phosphorylated form of eIF2α (Fig. 1A and B). Finally and despite the pTFEB increase, we observed an increase in the lysosomal marker LAMP1 protein 72h after the POS pulse (Fig 1A and B). Overall, these results show that upon feeding a large quantity of POS (*7*), RPE cells initially activate mTOR in the pulse period, followed by an mTOR inhibition during the chase period in order to boost their degradative capacity by increasing TFEB expression. Somehow, TFEB remains mostly inactive which prevents a more robust translation of lysosomal and autophagic components. These results raised the possibility that if cells could induce a stronger TFEB response, the result could be a more effective digestion of the phagocytosed material and a reduction of potentially toxic AFGs load.

### Inducing autophagy with rapamycin decreases POS-dependent autofluorescence through a cathepsin- and lipase-dependent mechanism

Rapamycin is the classical inhibitor of mTOR, specifically of mTORC1 (*25*). We confirmed that in our system, RPE cells incubated with rapamycin during 24h showed a reduction in pS6 (a readout of mTORC1 activity) starting at a concentration of 1 nM (Appendix Fig S2). Thus, hfRPE and ARPE-19 cells were pulsed with POS, and treated with rapamycin (10, 50 and 100 nM) starting at 48h after the POS pulse (Fig 2A). We decided to incubate RPE cells at this timepoint for several reasons; i) it is when POS-dependent AF is already detected (*7*), ii) the chase period between 24 and 72h seemed a good window of opportunity to act on TFEB, iii) it mimics the situation in AMD patients, where a significant load of lipofuscin granules are already established during early disease. We observed that rapamycin was able to significantly resolve AFGs accumulation reducing the total load of AF by about 30%, in both hfRPE and ARPE-19 cells (Fig. 2A). Confocal microscopy showed that the decrease in POS-dependent AF by rapamycin in ARPE-19 cells was associated with a significant decrease in the mean number of AFGs and not in the AFGs area (Fig. 2B).

**Fig. 2.**
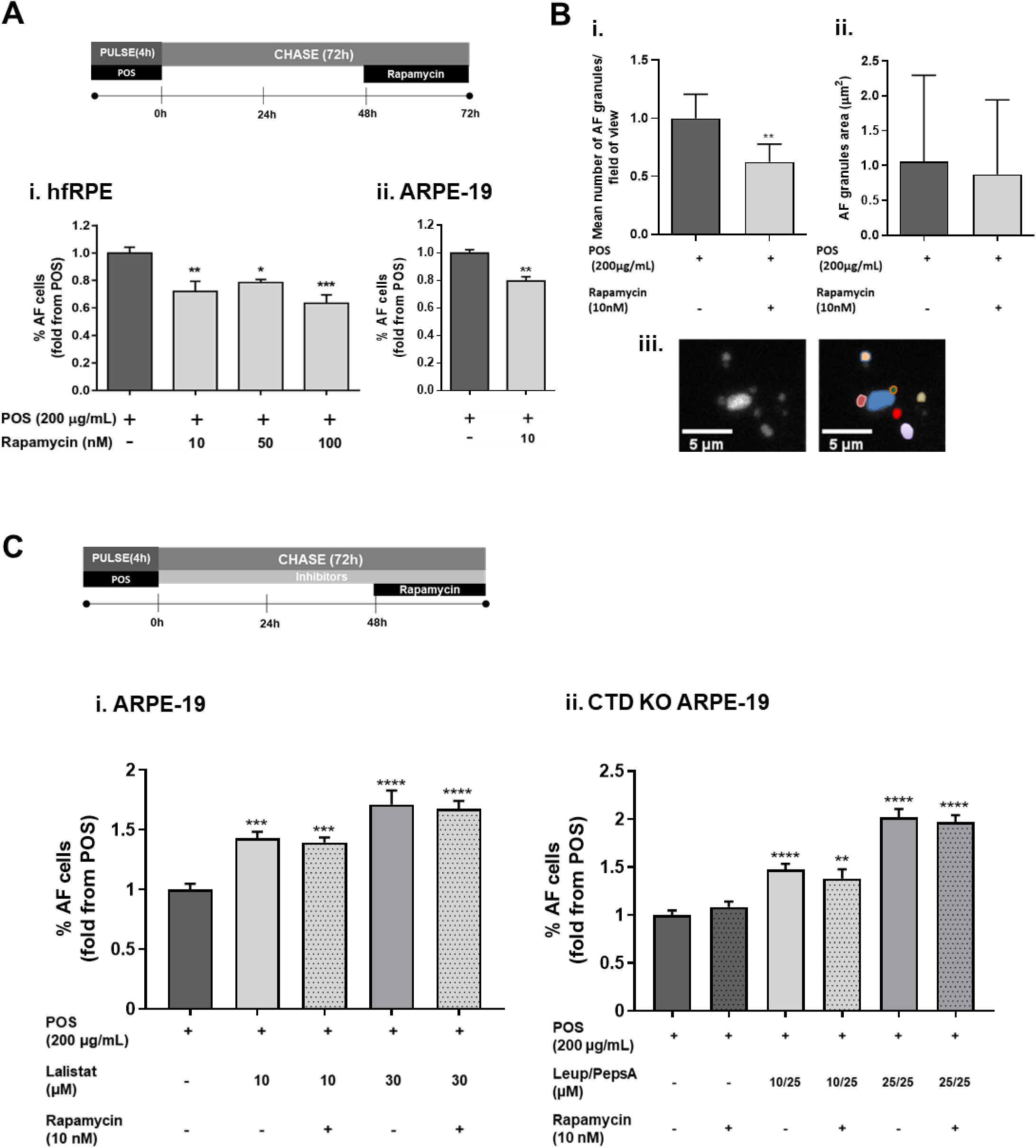
Effect of Rapamycin in POS-dependent autofluorescence. (A) RPE cells were pulsed during 4h with 200 μg/mL POS, washed and after 48h were incubated with rapamycin for 24h. The percentage of autofluorescent (AF) cells was evaluated by flow cytometry, as described in Materials and Methods. Results were expressed as fold change from POS values; (i) hfRPE were incubated with 10, 50 and 100 nM rapamycin and (ii) ARPE-19 cells were incubated with 10 nM rapamycin. (B) Microscopic analysis of the number and size of autofluorescent granules (AFGs) after a 72h chase period in ARPE-19 cells; (i) Quantification of the number of AFG per field of view (FOV), standardised to the mean of POS condition. (ii) Average size (area in μm2) of AFGs. (iii) Representative immunofluorescence image of five AFGs (left) and their automatic identification and segmentation (right) for number and size quantification (as described in Materials and Methods). Scale bar 5 μm. (C) Schematic of rapamycin treatment in the presence of lysosomal acid lipases or proteases inhibitors. Cells were pulsed 4h with 200 μg/mL POS and incubated for 72h with inhibitors and in the last 24h of the chase period rapamycin was added. The percentage of AF cells was evaluated by flow cytometry; (i) ARPE-19 cells were pulsed with 200 μg/mL POS and incubated for 72h with different concentrations of Lalistat2 (10 and 25 μM) and a 24h incubation with rapamycin (10 nM). (ii) ARPE-19 Cathepsin D KO cells were pulsed with 200 μg/mL POS and incubated for 72h with Leupeptin (10 and 25 μM) and pepstatin (25 μM) and a 24h incubation with rapamycin (10 nM). All data are shown as mean ± SEM of at least 3 independent experiments (n≥3). A one-way ANOVA followed by multiple comparisons Dunnet post hoc correction (A for hfRPE and C) and a Student’s t test was used for statistical analysis (A for ARPE-19 and B) and *P<0.05, **P<0.01, ***P<0.001 and ****P<0.0001 vs. POS (dark grey bars, no rapamycin).

We next explored the role of functional lysosomes in AFG degradation, by interfering with the hydrolytic activity of lysosomes during rapamycin treatment (Fig. 2C). In the presence of Lalistat-2, an inhibitor of lysosomal acid lipases the effect of rapamycin is neutralised. AF levels increase in a concentration-dependent manner regardless of the presence of rapamycin (Fig. 2C). We also used cathepsin D (CTSD) knock-out (KO) ARPE-19 cells as CTSD has been described as the primary lysosomal enzyme involved in POS degradation in RPE cells (*26, 27*). We have previously shown that CTSD KO ARPE-19 cells generate a higher load of AFGs when compared with non-mutated cells fed with the same amount of POS (*7*). Again, we observed that the rapamycin effect was neutralised in this mutated cell line (Fig. 2C). Finally, the use of lysosomal proteases inhibitors Leupeptin and Pepstatin A, further enhanced the formation of AF in a concentration-dependent manner and also neutralised the effect of rapamycin (Fig. 2C). These results suggest that the effect of rapamycin is dependent on functional lysosomes, both through protease and lipase-dependent hydrolysis.

To study alternatives to rapamycin and explore new therapeutic options, we analysed a collection of Boehringer Ingelheim’s known mTOR/autophagy modulators (Table 1). For this experiment, we used cross-linked uvPOS, which are described to be more difficult to digest (*28*), and the same incubation scheme as for rapamycin (Fig. 2A). From the tested compounds, INK-128 and compound H showed a significant decrease in POS-dependent AF (∼0.7-fold). These drugs have the same described mechanism of action as rapamycin, via mTOR inhibition.

**Table 1.** Effect of Boehringer Ingelheim’s drugs on UVPOS-dependent AF, as compared to Rapamycin.

| Drug name/Lab code | Mechanism of action | Concentration | POS-dependent AF (fold decrease from UVPOS±SEM) |
| --- | --- | --- | --- |
| INK 128 | mTOR inhibitor | 10nM | 0.72 ± 0.09* |
| B | Autophagy inducer, molecular target unknown | 100nM | 0.74 ± 0.09 |
| H | mTORC1 antagonist | 100nM | 0.67 ± 0.10* |
| I | mTORC1 antagonist | 10nM | 0.89 ± 0.08 |
| **Rapamycin** | **mTOR inhibitor, autophagy inducer** | **10 nM** | **0.61 ± 0.09\*\*** |

At least 3 independent experiments. \*P<0.05; \*\*P<0.01 vs. UVPOS-fed cells

### Rapamycin decreases POS-dependent autofluorescence by activating GADD34 downstream pathways and lysosomal biogenesis

We then explored the effect of rapamycin on the expression of TFEB and other lysosomal biogenesis-related proteins, using a time-course experiment after POS feeding (Fig 3A). In the presence of POS, and as expected, rapamycin significantly inhibited pS6, an indication of mTOR inhibition (Fig 3B and C). The pTFEB/TFEB ratio decreased with rapamycin treatment (Fig.3C) with a consequent increase in GADD34 protein. The GADD34 peak at 2h induces dephosphorylation of eIF2α, demonstrated by the significant decrease in peIF2α/eIF2α ratio 4h after rapamycin treatment. TFEB activation therefore leads to GADD34 activation and enables the transcriptional program for lysosomal biogenesis, which was corroborated by the increase in LAMP1 protein 24h after rapamycin treatment (Fig 3B, C and Fig 4). To further demonstrate the relevance of GADD34 in the normal POS degradation following phagocytosis by the RPE, we inhibited GADD34 with salubrinal (*24*). Upon salubrinal treatment, AFG levels increase significantly in ARPE-19 cells fed with POS (see Appendix Fig. S1).

**Fig. 3.**
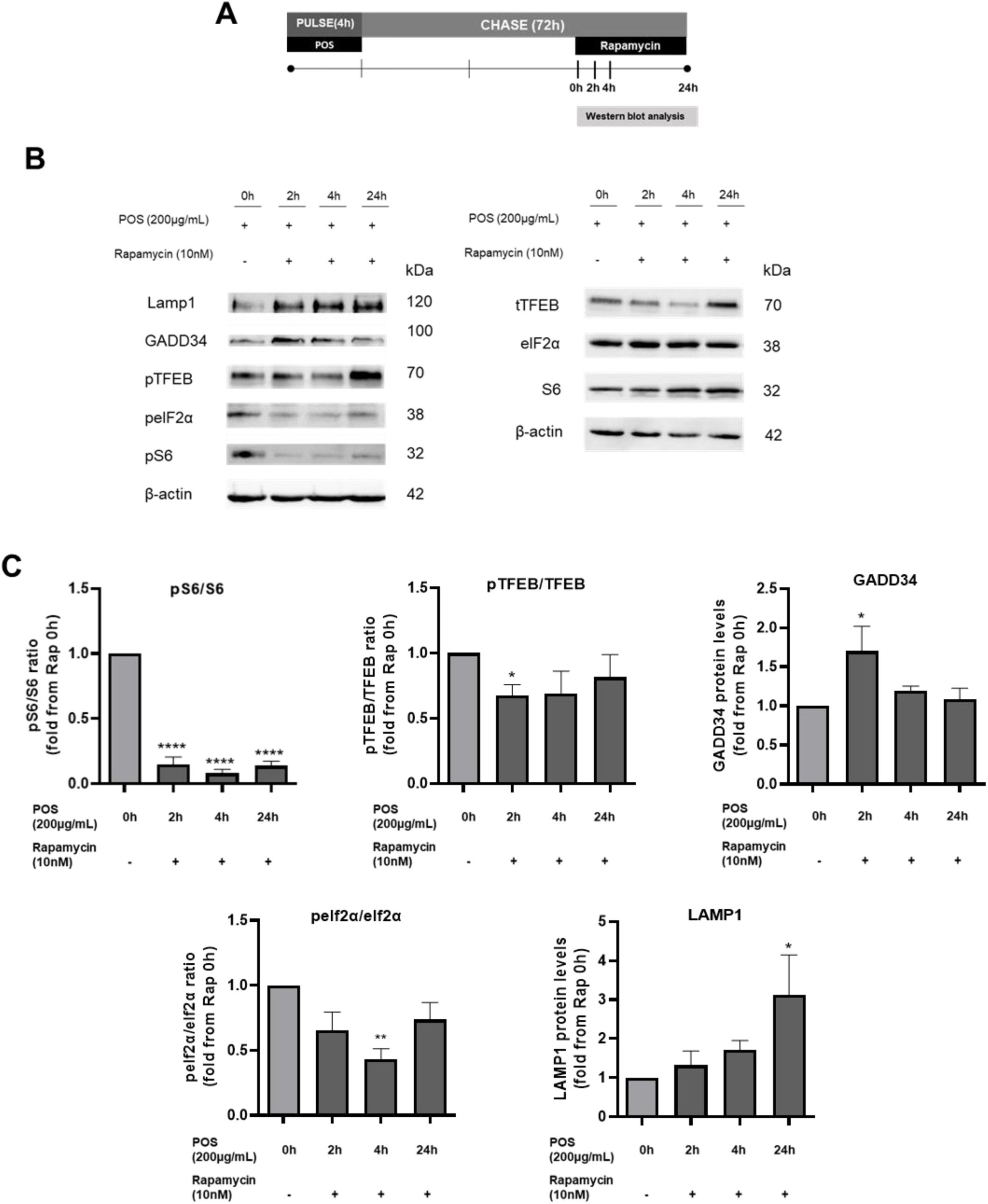
Rapamycin effect in TFEB and other lysosomal biogenesis-related proteins. (A) ARPE-19 cells were pulsed during 4h with 200 μg/mL POS, washed and after 48h were incubated with rapamycin (0h timepoint). (B) Evaluation of pS6, total S6, pTFEB, total TFEB, GADD34, peIf2α, total peIf2α, Lamp1 and β-actin protein levels at 2h, 4h and 24h after rapamycin incubation, assessed by western blot. Representative blots of one experiment are shown. (C) Densitometric analysis of the immunodetection of the indicated proteins was performed as described in Materials and Methods and the ratio of pTFEB/TFEB, pS6/S6 and peIf2α/eIf2α were calculated using β-actin as loading control. All data are shown as mean ± SEM of at least 3 independent experiments (n≥3). A one-way ANOVA followed by multiple comparisons Dunnet post hoc correction was used for statistical analysis and *P<0.05, **P<0.01 and ****P<0.0001 vs. POS 0h (grey bars, no rapamycin).

**Fig. 4.**
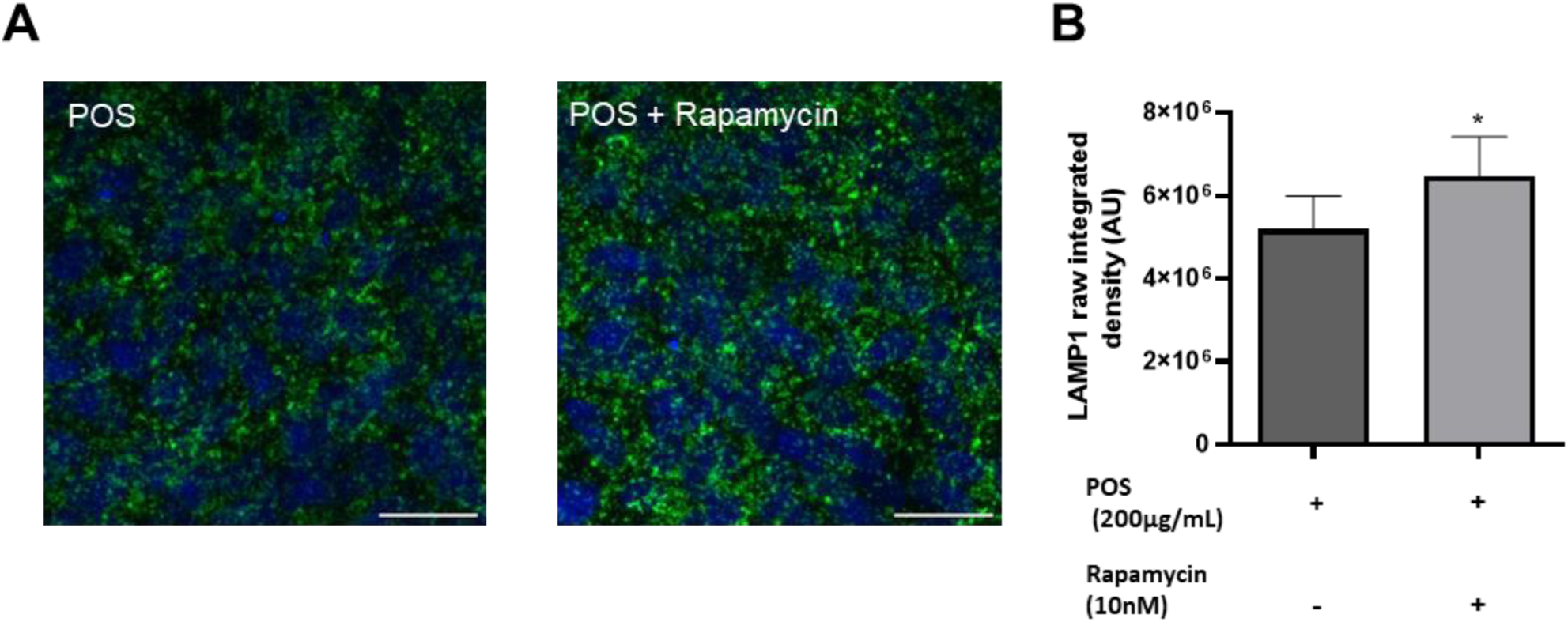
Effect of Rapamycin on LAMP1 expression. ARPE-19 cells were pulsed during 4h with 200 μg/mL POS, washed and after 48h were incubated with rapamycin during 24h. (A) After incubation, cells were fixed and stained with mouse anti-LAMP1 (green) and DAPI (blue), as described in Materials and Methods. Representative images of a Z-slice are shown. Scale bar: 30 µm. (B) LAMP1 integrated density was quantified using ImageJ and data are shown as mean ± SEM at least 3 independent experiments (n≥3). A Student’s t test was used for statistical analysis and *P<0.05 vs. POS (dark grey bars, no rapamycin).

These results confirm our hypothesis that activation of TFEB through mTOR inhibition can improve clearance of lipofuscin-like AFGs in RPE cells. Moreover, they point to the relevant role of the downstream target of TFEB, GADD34, in this process.

### Genetic targeting of TFEB can potentially be used to reduce POS- dependent autofluorescent granule load

As a proof of concept, we next investigated whether inducing TFEB expression and/or its nuclear translocation could prevent and/or partially resolve POS- dependent AF. To test this hypothesis, we transduced hfRPE with AAVs to overexpress TFEB or a constitutively active (CA) form of TFEB (TFEB-CA). TFEB-CA (TFEB^S142A,S211A^) is constitutively active as it cannot be phosphorylated by mTOR on serine residues S142 and S211 (*20, 29*). Transcriptional and protein levels of TFEB were assessed as a proof of RPE successful transduction (Appendix Fig S3). Cells were transduced immediately after the POS pulse mimicking the rapamycin protocol (resolving approach, Fig. 5A). A significant decrease in AFGs was observed in cells overexpressing the TFEB-CA form. When RPE cells were transduced days before POS feeding (preventive approach, Fig. 5B), we observed a striking reduction in the appearance of POS- dependent AFGs, when TFEB is constitutively active. These results further demonstrate that strategies modulating TFEB activation to improve lysosomal function can potentially be used not only as a therapeutic approach in AMD but also in diseases where the accumulation of undigested toxic material causes cellular dysfunction.

**Fig. 5.**
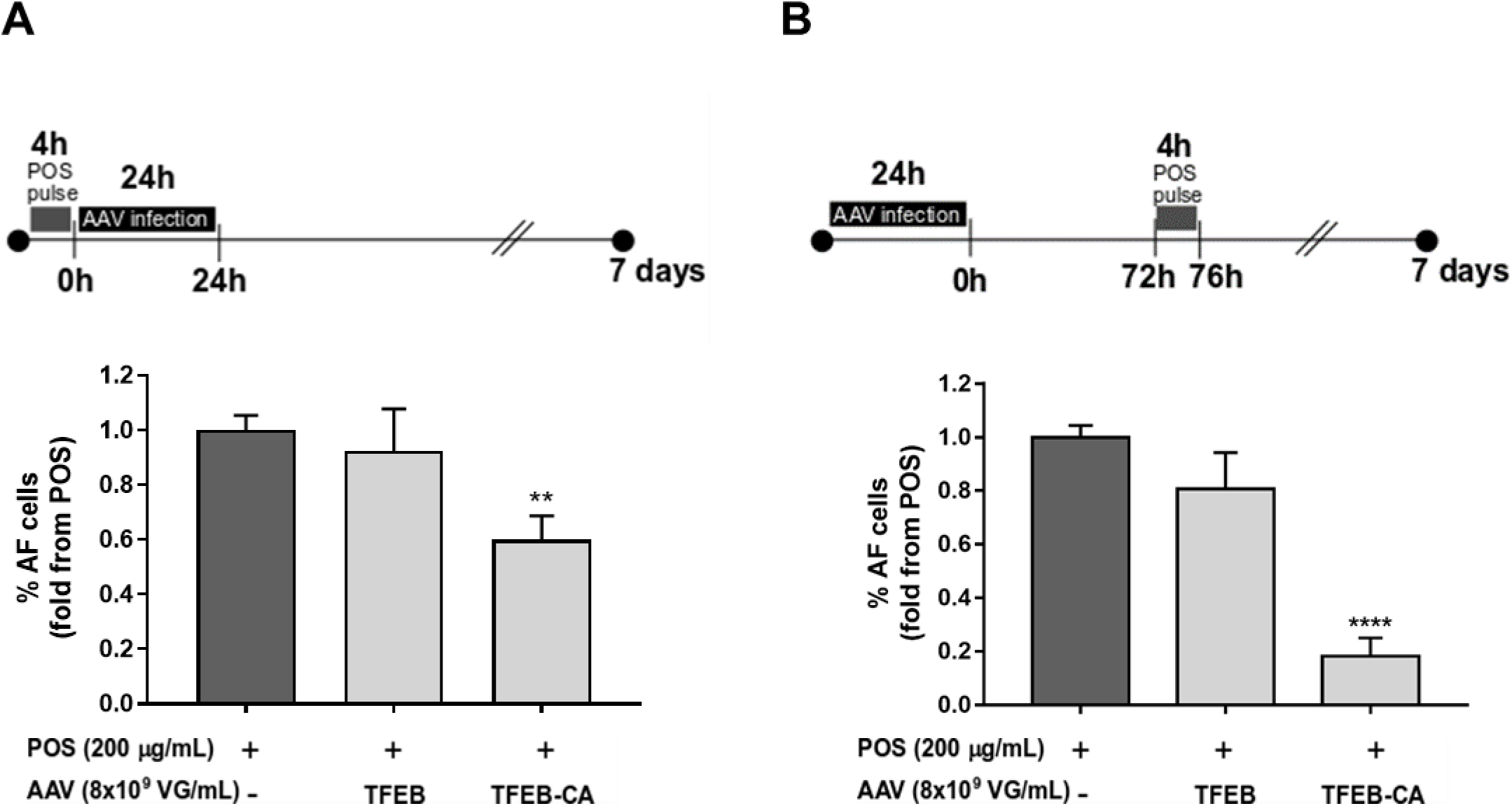
Effect of Adenovirus-associated viral vectors (AAVs) therapeutic strategies in POS-dependent AF. (A) As a resolving strategy, hfRPE cells were first pulsed with POS for 4h and then infected with the AAVs, with a chase period until 7 days. (B) A preventive approach was tested by transducing hfRPE cells for 24h hours prior to POS pulse (4h), followed by a 72h chase period. The percentage of autofluorescent (AF) cells was evaluated by flow cytometry, as described in Materials and Methods. Results were expressed as fold change from POS values. All data are shown as mean ± SEM of at least 3 independent experiments (n≥3). A one-way ANOVA followed by multiple comparisons Dunnet post hoc correction was used for statistical analysis and **P<0.01 and ****P<0.0001 vs. non-infected POS-fed cells (dark grey bars).

## DISCUSSION

The work presented here suggests that targeting the TFEB/mTOR axis could potentially reduce lipofuscin build-up in RPE cells normally found during aging and exacerbated in AMD. Robust TFEB activation leads to lysosome biogenesis and new lysosomes clear a significant proportion of the accumulated AFGs made of undigested POS (Fig 6). As the accumulation of AF material (lipofuscin) *in vivo* is potentially toxic, modulating the mTOR/TFEB axis could be beneficial in the early/intermediate stages of AMD to delay disease progression.

**Fig. 6.**
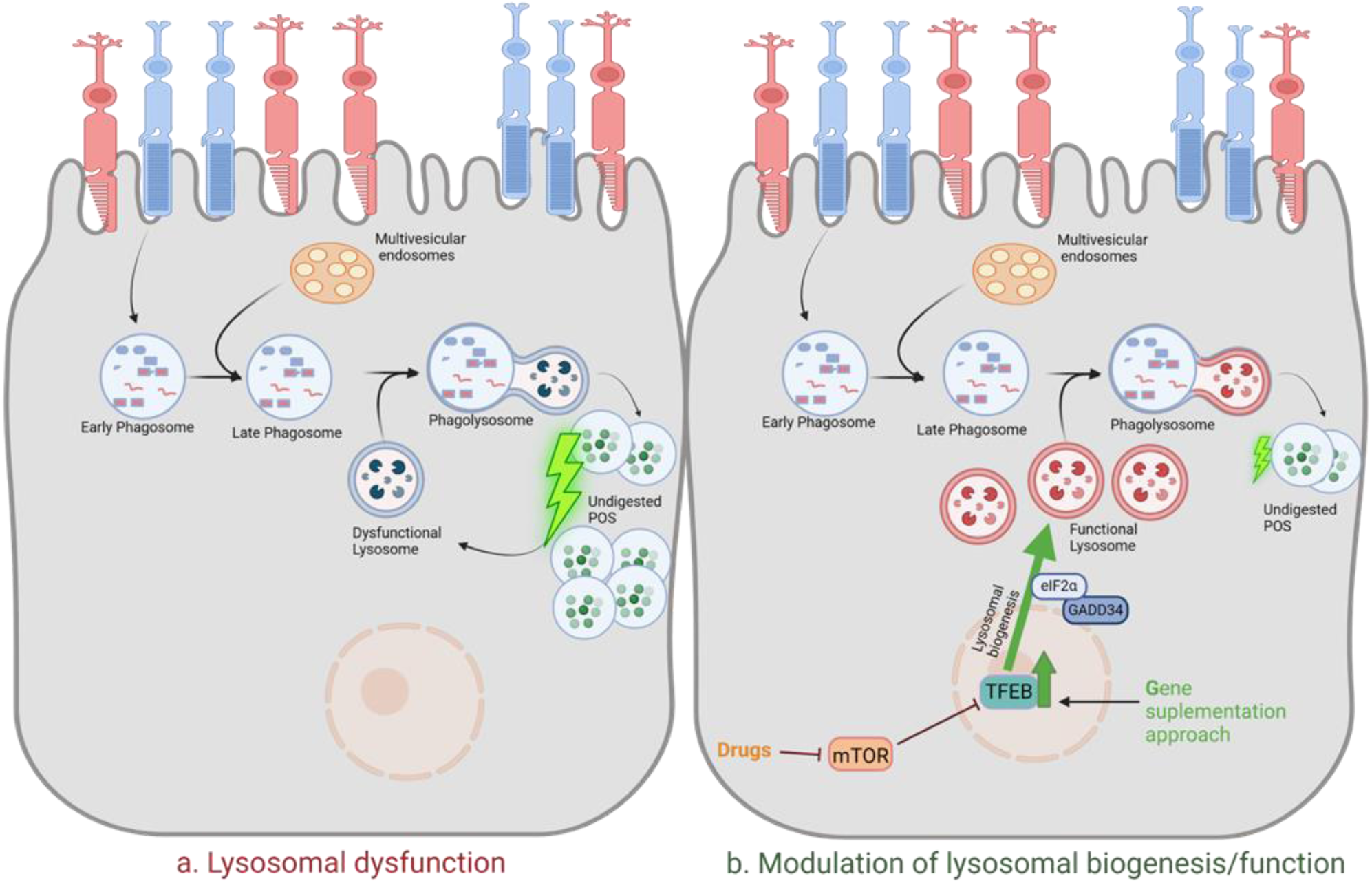
Proposed model for modulation of the autofluorescent granule load in RPE cells. (A) For AFG formation, POS are phagocytosed by the RPE. Early phagosomes can interact with multivesicular endosomes forming late phagosomes that will fuse with lysosomes originating phagolysosomes. When cells are unable to degrade the POS load, the accumulation of undigested POS leads to lysosomal dysfunction, further increasing the amount of undigested material and resulting in POS-dependent autofluorescence. (B) We propose a therapeutic strategy to decrease the AFG load by targeting the mTOR/TFEB axis. We propose to specifically inhibit mTOR or activate the constitutively active form of TFEB, which will increase lysosomal biogenesis, a process that is mediated by GADD34 through eIF2α dephosphorilation, ultimately leading to an increase in the number of functional lysosomes, which in turn will will decrease the AFG load in RPE cells. Created with BioRender.com.

The molecular mechanisms responsible for the phagocytosis and clearance of POS have yet to be fully understood. Cargo degradation by both phagosomes and autophagosomes is sensitive to lysosomal biogenesis, and changes in the levels of lysosomal proteins have been linked to various retinal diseases (*30–32*). Here we observed mTOR transiently activated upon POS phagocytosis. This observation agrees with *in vivo* studies in mouse RPE/choroid whole-mounts following light onset, a trigger of POS phagocytosis (*23*). We asked whether the mTOR activation was sustained and showed that mTOR was inhibited after 8h post-phagocytosis *in vitro*. Accordingly, we observed gradually increasing levels of TFEB over the course of the chase period, however surprisingly TFEB remains largely inactive, suggesting a bottleneck in achieving a robust lysosomal and autophagic activation. Corroborating the present results, an increased phosphorylation of TFEB was observed in RPE lysates from human AMD donors but not in age-matched non-AMD controls (*33*).

TFEB is a master regulator of autophagy and lysosomal biogenesis (*34*). TFEB can be regulated by mTORC1, which in turn regulates lysosomal activity by directly preventing autophagy and TFEB activation. In aged RPE cells, increased mTORC1 activity inhibits POS degradation and may further exacerbate lysosome dysfunction (*35*). Importantly, decreased autophagy flux was also observed on the RPE from AMD donors (*12*). Accordingly, the pharmacological inhibition of mTOR emerges as a very attractive approach. Indeed we show here that rapamycin, a thoroughly studied mTORC1 inhibitor (*25*), significantly reduces POS-dependent AF formation in RPE cells by enhancing lysosomal biogenesis. Rapamycin treatment led to the activation of GADD34, an eIF2α phosphatase and TFEB target, which is crucial for enhancing lysosomal biogenesis (*24*). Correspondingly, inhibition of GADD34 with salubrinal further increased AFG accumulation in our assay. Furthermore, we show a significant increase in LAMP1, a marker of the lysosomal mass in cells. This indicates that mTOR inhibition facilitates TFEB activation, which in turn promotes the degradation of phagocytosed material and reduces AFG accumulation Finally, we show that the effect of rapamycin is dependent on active lysosomal proteases and lipases, highlighting the necessity of functional lysosomes for effective clearance of AF material.

The effect of rapamycin in decreasing AF in RPE cells has been studied previously in other contexts. In ARPE-19 cells treated with A2E, a major component of toxic lipofuscin implicated in AMD, rapamycin was described to stimulate autophagy via the inhibition of mTOR, reducing A2E accumulation and attenuating inflammation-associated and angiogenic factors (*36*). Moreover, ARPE-19 cells treated with rapamycin and fed with HNE-modified POS result in a decrease in lipofuscin-like autofluorescence (*37*). Additionally, rapamycin also induces a significant accumulation of TFEB in the nucleus of RPE cells (*38*). Finally, in a sodium-iodate (NaIO3)-induced retinal degeneration mice model, often used as a model of AMD, rapamycin played a key role in attenuating an inflammatory response and oxidative stress (*39*).

Future directions point to clinical translation and/or combination therapies. Nevertheless, the potential use of rapamycin to treat AMD should be taken very carefully. While rapamycin or rapalogs (*40*) have shown therapeutic efficacy for age-related diseases in animal models, significant side effects are restraining their use in humans. Recent studies suggest synthetic high-density lipoprotein nanoparticles delivering rapamycin as a more efficient therapeutic delivery option to treat AMD (*41*). The demand for effective disease-modifying strategies against age-related diseases is currently stimulating the development of a wide number of novel molecules, as well as the re-evaluation of old drugs for their pro-autophagic potential. With this in mind, we tested four Boehringer Ingelheim proprietary molecules recognized as autophagy inducers but some with still unknown molecular targets. Although the majority of compounds were able to decrease the AF induced by uvPOS feeding, this effect was only significant with INK 128 and Drug H, both of which act via mTOR, supporting its central role in this process (*42*).

As a proof-of-concept to demonstrate the critical role of TFEB in the clearance of AFGs, we show that overexpression of a constitutively active form of TFEB (TFEB-CA) significantly reduced POS-dependent AF accumulation in RPE cells, both in preventive and resolving approaches. This demonstrates the therapeutic potential of strategies aimed at activating TFEB to enhance cellular clearance mechanisms. However, the use of gene therapy to constitutively express TFEB seems ill-advised as rare kidney cancers are caused by this mechanism (*15*). Thus, we suggest investigating the combination of TFEB activation with other therapeutic strategies, such as anti-inflammatory and/or antioxidant treatments, which could provide a more comprehensive approach to managing AMD.

In summary, our study provides compelling evidence that modulating the TFEB/mTOR axis can significantly improve lysosomal function and reduce the accumulation of toxic AF material in RPE cells. Our results underscore the therapeutic promise of TFEB activation and mTOR inhibition as strategies to combat the progression of AMD and preserve vision in the aging population.

## MATERIALS AND METHODS

### Cell Cultures

Primary human fetal RPE cells (hfRPE) were purchased from Lonza and cultured using optimized RtEBM^TM^ Basal Medium and RtEGM^TM^ SingleQuots^TM^ Supplements (Lonza), as already described (*7*). The ARPE-19 cells, a human RPE cell line (ATCC), were cultured in Dulbecco’s Modified Eagle Medium/Nutrient Mixture F-12 (DMEM/F-12) (Gibco) supplemented with 10% fetal bovine serum (FBS) (Gibco) and 1% Penicillin-Streptomycin (Pen/Strep) (Gibco). Cathepsin D (CTSD) knock-out (KO) ARPE-19 cells were generated as previously described (*7*). All of the cell models were grown in a 5% CO2 incubator at 37 °C.

### POS Phagocytosis Assays

Photoreceptor outer segments (POS) were isolated from porcine eyes, according to Parinot et al. (*43*), with some minor modifications (*7*). Before the phagocytosis assays, POS preps were thawed in the dark, washed three times in PBS by spinning for 10 min at 2400 xg. For the UV-irradiated POS (uvPOS), each prep was exposed to ultraviolet radiation using a UVP Crosslinker (CL-1000 Model, Analyticjena) with 2×2 minutes pulses of 254 nm at an estimated radiant exposure of 1 J/cm^2^.

RPE cells were fed during 4h (pulse period) with POS at a final concentration of 200 µg/mL (∼2.6x10^5^ POS particles/cm^2^ or ∼1x10^6^ POS particles/mL) in DMEM/F- 12 supplemented with 10% FBS. After this pulse, POS were removed, cells were washed and cultured in supplemented DMEM with 1% FBS for hfRPE or with 10% FBS for ARPE-19 cells (chase period). To study the mechanisms of POS phagocytosis and clearance, the beginning of feeding was considered as 0h. Cells were trypsinized for Western blot analysis during the pulse (1h, 2h and 4h) and chase (8h, 24h, 72h and 7 days) periods and for quantitative real-time-PCR during the chase (24h and 72h) period.

### Drug treatments

After 4h of POS feeding, hfRPE cells were washed and cultured in supplemented DMEM with 1% FBS during 48h and then incubated during the last 24h of the chase period with rapamycin (10, 50 and 100 nM; Sigma-Aldrich). At the end of the incubation, cells were trypsinized for flow cytometry to analyse the AF. In ARPE-19 cells, after the 4h POS feeding, cells were washed and cultured in supplemented DMEM with 10% FBS during 48h and then incubated during the last 24h of the chase period with rapamycin (10 nM) and then trypsinized for flow cytometry to measure AF or analysed by confocal microscopy, to assess the mean number of AFG/field of view, as well as the AFG area.

ARPE-19 cells were also incubated with Salubrinal (10 and 25 μM), a GADD34 inhibitor (*24*) during the 72 h chase period and then cells were trypsinized for flow cytometry and AF was measured.

To study the potential mechanisms of action of rapamycin in POS clearance, the beginning of rapamycin treatment (10 nM) in ARPE-19 cells was considered as 0h and cells were trypsinized for Western blot analysis after 2h, 4h and 24h.

To study the role of lysosomal function in the rapamycin effect, we used two different experimental approaches, after 4 hours of POS pulse: i) ARPE-19 cells were treated during the 72h chase period with Lalistat-2, a lysosomal acid lipase inhibitor (10 and 30 µM; Sigma-Aldrich) with or without rapamycin 10 nM and ii)

CTSD KO ARPE-19 cells were treated during the 72h chase period with the following lysosomal acid protease inhibitors Leupeptin (10 or 25 µM; Calbiochem, San Diego, CA) plus Pepstatin A (25 µM; Sigma-Aldrich) with or without rapamycin 10 nM. After the chase period, the cells were trypsinized for flow cytometry and AF assessed.

In another set of experiments, hfRPE cells were fed with uvPOS during 4h, washed, and cultured in supplemented DMEM-F12 with 1% FBS during 48h adding rapamycin (10 nM), or other potential autophagy inducers, provided by Boehringer Ingelheim (Table 1) in the last 24h of the chase period.

### AAVs transduction

hfRPE were transduced with Adeno-associated virus serotype 2 (AAV-2) to overexpress TFEB, a constitutively active (CA) form of TFEB (TFEB-CA (TFEB^S142A-S211A^)), (all constructed were provided by Boehringer Ingelheim), at a MOI of 2×10^9^ VG/well in a 48-well plate with supplemented DMEM with 1% FBS during 24h. Transcriptional and transductional expression was confirmed at 24h, 72h and 7days post-infection.

Two different experimental approaches were tested: a) cells were transduced during 24h and 72h later were fed with POS (4h), followed by a 72h chase period; and b) cells were fed with a POS (4h) and then transduced with the AAVs during 24h, with a total chase period of 7 days. At the end of both approaches, cells were trypsinized for flow cytometry.

### Flow Cytometry

For flow cytometry assays, cells were trypsinized with TrypLE™ Express Enzyme (Gibco) for 30 min. Cells were resuspended in DMEM supplemented with 10% FBS, washed two times with PBS and one time with flow cytometry buffer (1% FBS and 2 mM EDTA in PBS). Finally, cells were resuspended in flow cytometry buffer and data acquisition was performed in a FACS CANTO II flow cytometer (BDBiosiences) using the 488 nm excitation wavelength to evaluate cellular AF. At least 30,000 cells were acquired per condition using BD FACSDivaTM software (Version 6.1.3, BD Biosciences). Data analysis was performed in FlowJo (Version 10, BD Biosciences). Results from independent experiments, performed in triplicate, were represented as the percentage of 488-positive cells and were normalized to the values of AF detected in cells pulsed with POS but in the absence of compounds. Fold values were used for statistical analysis.

### Western Blot

Cells were lysed in ice cold cell lysis buffer (Cell Signaling Technology) supplemented with protease and phosphatase inhibitor cocktails (Roche) according to the manufacturer’s instructions. Lysates were pelleted for 15 min at 13 000 xg at 4 °C and supernatants kept for protein quantification (BCA Protein Assay Kit, Thermo Scientific). Equal amounts of cellular proteins were resolved on 12% sodium dodecyl sulfate – polyacrylamide gels (SDS-PAGE) and subsequently transferred to nitrocellulose membranes (Bio-Rad Laboratories). Membranes were blocked using 5% non-fat dry milk or 5% bovine serum albumin (BSA) (Sigma-Aldrich) for phosphorylated proteins immunoblots in Tris-buffered saline (TBS) (50 mM Tris, 150 mM NaCl, pH = 7.6) containing 0.1% Tween-20 (Sigma-Aldrich) (TBS-T) for 1 h. Primary antibody incubations were carried out at 4°C overnight (Appendix Table S1). After washing with TBS-T, the appropriate HRP-conjugated secondary antibody was added (1:5000 in blocking buffer) for 2 h at room temperature. Antibody binding was detected using chemiluminescence ECL Prime Western Blotting Substrate (GE Healthcare) and a ChemiDoc Touch Imagine System (Bio-Rad). We analyzed the band intensity using ImageJ analysis software (NIH), using β-actin as loading control. The intensity value was converted to fold change in comparison with control group (no POS). Fold values were used for statistical analysis.

### Quantitative real-time-PCR (qPCR)

RNA was extracted using the RNAeasy extraction kit (Qiagen) according to the manufacturer’s instructions. cDNA was prepared using Maxima first strand cDNA synthesis kit (ThermoFisher Scientific). Real-time quantitative polymerase chain reaction (RT-qPCR) was performed on a QuantStudio™ 5 (Applied Biosystems) using iTaq SYBR Green Supermix (ThermoFisher Scientific). The reaction mixture without template cDNA was run as a control and 3 replicates from each sample were performed. mRNA expression levels were normalized to HRPT1 and/or PGK1 house keeping genes, as indicated in Figure legends and relative gene expression levels were analysed using the Comparative CT Method (2-ΔΔCt method). Fold values were used for statistical analysis. Primer sequences used for PCR are listed in Appendix Table S2.

### Confocal Immunofluorescence Microscopy

Cells grown on coverslips were fixed for 15 min in 4% paraformaldehyde (Alfa Aesar) at room temperature. Cells were blocked/permeabilized for 30 min in PBS containing 1% BSA and 0.05% saponin or 0.1% TX100, according to the antibodies used. AF was visualized in the 405 nm excitation wavelength. For LAMP1 staining, cells were incubated with a mouse anti-LAMP1 conjugated with Alexa fluor 647 (BioLegend, clone H4A3, Cat #328609, 1:500) for 1h. Cell nuclei were labelled with DAPI (Sigma) (1 µg/ml) and cells were mounted in Mowiol mounting media (Calbiochem). Images were acquired in a Zeiss LSM 710 confocal microscope or in a Zeiss LSM980 airyscan confocal in the Multiplex SR- 4Y imaging mode, with a Plan-Apochromat 63x1.4 NA oil-immersion objective. All images within the same experiment were acquired with the same acquisition settings. Digital images were analysed using ImageJ (https://imagej.nih.gov/ij/). The number and area of AFG per field of view were automatically segmented and quantified using ImageJ (ver.2.9) (minimum of 8 field of view per experiment). Results from AFG analysis were represented as the mean number of AFG detected per field of view ± SEM. Lamp1 staining was quantified by choosing a manual threshold and the total raw integrated density was quantified per field of view (minimum of 5 field of view per experiment). Results were represented as mean integrated density ± SEM.

### Statistics

Statistical analysis was performed using GraphPad Prism, version 9 Software. All results are shown as mean ± SEM. Normal distribution of data was assessed using the Shapiro-Wilk normality test. Data that followed a normal distribution was analysed using Student t test or one-way ANOVA followed by multiple comparisons Dunnet post hoc correction, as appropriate. Data that did not follow a normal distribution was analysed using one-way ANOVA Kruskal Wallis test. p values less than 0.05 were considered statistically significant.

## Acknowledgments

We would like to acknowledge the Cell Tissue Culture, Flow cytometry and Microscopy facilities at NMS, as well as CASO (Centro de Abate de Suínos do Oeste) that generously provided porcine eyes for POS isolations.

## Author contributions

MCS, ST, TC, PN and ASF conceived and coordinated the study. ASF, CE, MLS, MP, IPS, RC and ISF performed the experiments. PA provided support in data analysis and interpretation. ASF and MCS wrote the original manuscript draft. CE, PA, ST, and MLS reviewed the manuscript. All authors approved the results and final version of the manuscript.

## Data and materials availability

All data needed to evaluate the conclusions in the paper are present in the paper or in the Supplementary Materials.

**Correspondence** and request for materials should be addressed to MCS

