## Supplementary material for "Enhancing lysosome function via mTOR/TFEB activation reduces lipofuscin-like granules in early Age-related Macular Degeneration": Table S1; Table S2; Appendix Fig. S1; Appendix Fig. S2; Appendix Fig. S3

### Supplementary Materials

**Table S1**. List of primary antibodies used for Western blot

| Primary Antibody | Dilution (reference; brand) |
|---|---|
| **TFEB** | 1:1000 (#4240, Cell Signaling) |
| **pTFEB (Ser211)** | 1:1000 (#37681, Cell Signaling) |
| **GADD34** | 1:1000 (#10449-1-AP, Proteintech) |
| **eIF2α** | 1:1000 (#5324T, Cell Signaling) |
| **peIF2α (Ser51)** | 1:1000 (#9721S, Cell Signaling) |
| **S6** | 1:1000 (#2217S, Cell Signaling) |
| **pS6 (Ser 235/236)** | 1:1000 (#4858S, Cell Signaling) |
| **Lamp1** | 1:1500 (#3243S, Biolegend) |
| **V5** | 1:1000 (sc-83849, Santa Cruz) |
| **β-actin-peroxidase** | 1:25000 (A3854, Sigma) |

**Table S2**. List of primer sequences used for qPCR

| Gene | Oligonucleotides |
|---|---|
| **TFEB** | F: 5’- AGCTCACAGATGCTGAGAG -3’ |
| | R: 5’- TGTTGAACCTTCGTCTCCT -3’ |
| **GADD34** | F: 5’-TGAGACTCCCCTAAAGGCCA -3’ |
| | R: 5’ -CCAGACAGCCAGGAAATGGA -3’ |
| **LAMP1** | F: 5’- CGTGTCACGAAGGCGTTTTCAG -3’ |
| | R: 5’ - CTGTTCTCGTCCAGCAGACACT-3’ |
| **TFEB (exogenous)** | F: 5’- GAACAAGGGCACCATCCTGA -3’ |
| | R: 5’- GCCGTCTGCTGTGATTTTCC -3’ |
| **HPRT1** | F: 5’- GACCAGTCAACAGGGGACAT -3’ |
| | R: 5’- CCTGACCAAGGAAAGCAAAG-3’ |
| **PGK1** | F:5’-AACAACCAGAGGATTAAGGC-3’ |
| | R:5’-GCTCATAAGGACTACCGAC-3’ |

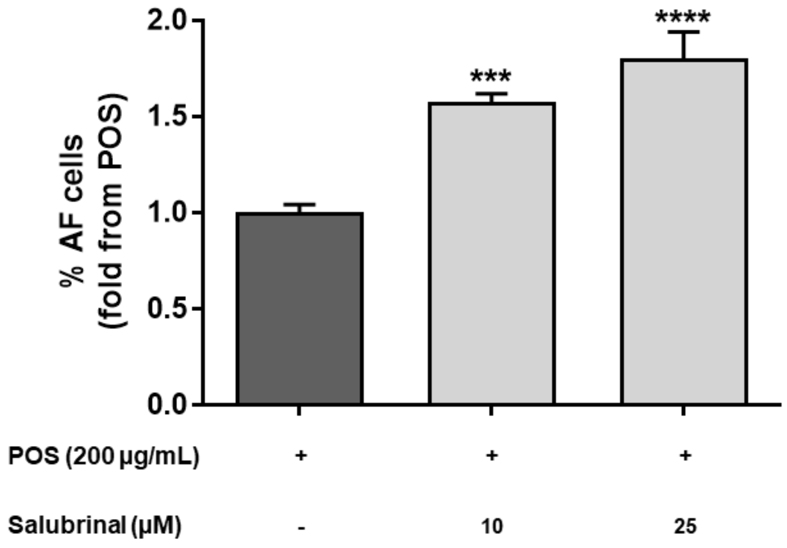


**Appendix Fig. S1- Salubrinal effect in POS-dependent autofluorescence**. ARPE-19 cells were pulsed with 200 µg/mL POS and incubated for 72h with salubrinal (10 and 25 µM). The percentage of autofluorescent (AF) cells was evaluated by flow cytometry, as described in Materials and Methods and results were expressed as fold change from POS values. All data are shown as mean ± SEM of at least 3 independent experiments (n≥3). A one-way ANOVA followed by multiple comparisons Dunnet post hoc was used for statistical analysis and ***P<0.001 and ***P<0.0001 vs. POS (dark grey bars, no salubrinal).

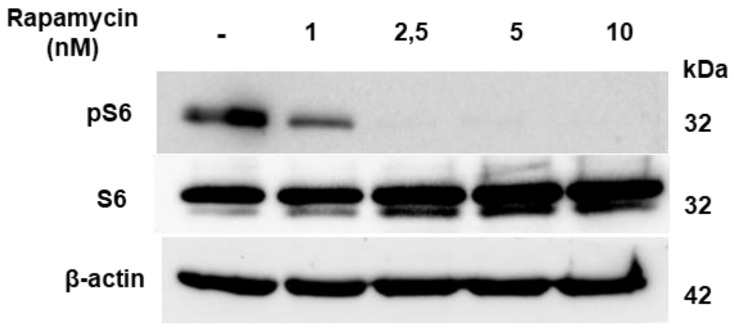


**Appendix Fig. S2- Inhibition of PS6 by rapamycin.** ARPE-19 cells were incubated for 24h days with different concentrations of Rapamycin (0, 1, 2,5, 5 and 10 nM). Evaluation of pS6, total S6 and β-actin protein levels, assessed by western blot. Representative blot of one experiment is shown.

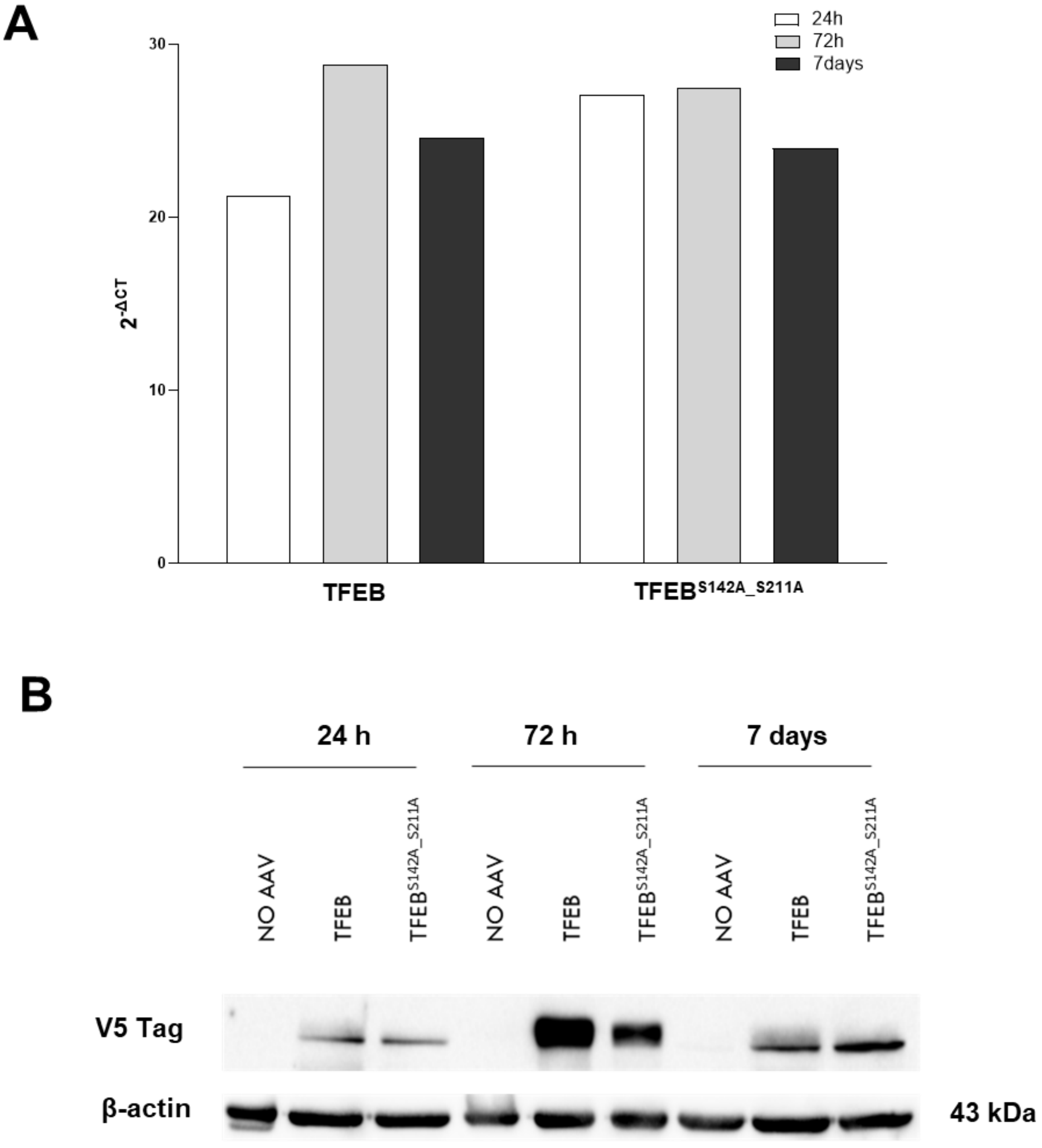


**Appendix Fig. S3- Transcriptional and protein levels of both TFEB forms in infected hfRPE as a proof of AAV infection.** (A) Exogenous mRNA levels of TFEB and TFEB $^{S142A\_S211A}$ were assessed by quantitative real time-PCR using HRPT1 and PGK1 as housekeeping genes. (B) The translated proteins were detected by western blot, using an antibody against V5 (specific tag for efficiently infected cells) and no infected cells as negative controls. Representative blot of one experiment is shown.
